# Structure and function of human NXPE1, a sialic acid *O*-acetyltransferase

**DOI:** 10.64898/2026.05.20.726592

**Authors:** H Zhang, F Li, M Zhang, Hiroyasu Konno, Wei Yu, X Min, Mousumi Paulchakrabarti, B Choudhury, Antony Simons, DE Piper, H Hsu, W Ouyang

## Abstract

Human genetic studies have identified defects in multiple mechanisms that predispose the risk of developing inflammatory bowel diseases (IBD), which include alterations in adaptive and innate immune responses, epithelial integrity and regulation of the intestinal mucus layer. Despite the importance of intestinal barrier integrity in the pathogenesis of IBD, essentially all current therapies modulate the immune responses. In this study, we determined the high resolution cryo-EM structure of human NXPE1, a IBD associated protein. Based on the structural homology, we identified NXPE1 as an *O*-acetyltransferase. Since NXPE1 is a *pseudo* gene in mouse, we generated knockout mouse model that lacked two of the mouse NXPE1 homologs, Nxpe2 and Nxpe4. The *O*-acetylation of sialic acid on red blood cells was abolished in the double knockout mice, confirming the sialic acid *O*-acetyltransferase function of NXPE1 family members. These findings underscore the potential of NXPE1 as a novel therapeutic target of the intestinal barrier functions for the treatment of IBD.

## Introduction

Inflammatory bowel disease (IBD) is classified into two major forms, ulcerative colitis (UC) and Crohn’s disease (CD), which are progressive chronic inflammatory conditions affecting the barrier integrity and normal functions of the intestinal tract. UC and CD have clinical, endoscopic and histopathological differences [1]. The exact causes of IBD remain elusive.

Genetic predisposition, environmental factors, change of gut microbiota, and loss of immune tolerance all contribute to the initiation of IBD. The onset of IBD is usually associated with compromised barrier functions and uncontrolled immune cell activation, which can form a vicious loop promoting further tissue damage and exacerbation of the inflammatory responses[2].

Genome wide association studies have identified more than 240 single nucleotide polymorphisms (SNPs) that are linked to the risk of development of IBD[3]. These genetic associations provide important insights into the pathogenesis of IBD. Epithelial integrity and homeostasis, microbial sensing and defense, innate and adaptive immune pathways have all been identified by GWAS to be associated with IBD. Autophagy, ER stress, cell migration, apoptosis/necroptosis, carbohydrate metabolism and oxidative stress are pathways impacting barrier functions and innate host defense[4-8]. Despite the complexity in the cellular and genetic pathways associated with the pathogenesis of IBD, the current standard treatments for IBD primarily target immune related mechanisms [9-12]. Importantly, only a small percentage of patients can achieve remission under these treatments. New therapies modulating other aspects of IBD pathogenesis may be necessary to help a broader population of patients.

A healthy intestinal barrier consists of intestinal epithelial cells with a mucus layer on the top. The intestinal barrier not only functions in nutrient and water absorption, but also sequesters luminal commensals and pathogenic microbes as well as their antigens from directly interacting with intraepithelial lymphocytes (IELs) and lamina propria immune cells[13]. In addition, the presence of various immune repressive cells such as T regulatory cells (Tregs) preserve the inner lamina propria in a non-inflamed state [14]. However, a breached intestinal barrier layer increases the infiltration of pathogens and antigens into lamina propria, which breaks immune tolerance and triggers inflammatory cascades. If the condition is not reversed back to intestinal homeostasis, it may evolve to chronic relapsing IBD[14]. There are two mucus layers in the colon, forming the essential barriers to protect the integrity of the intestinal outlining [15, 16]. The out-layer form is loose and mobile, which interacts and shapes the intestinal microbiome. The inner layer is firmly attached to the epithelial cells, dense and free of bacteria under homeostatic conditions. However, it becomes penetrable with microbes in patients with UC and in mouse models of colitis [17]. Muc2 is one of the major components of the colon mucus layer and predominantly produced by goblet cells [18]. The essential role of Muc2 and the mucus layer is revealed in Muc2 deficient mice, which develop colitis spontaneously as early as 5 weeks of age, and intestinal tumors[19, 20]. The epithelial barrier from Muc2 deficiency is associated with abnormal crypt morphology, mucosal thickening, directly contacting with microbes, and immune infiltration and activation.

Muc2 is highly glycosylated with mucin-type *O*-linked oligosaccharides (*O*-glycans), the major type glycan in the gut[21]. Epithelial glycans provide bacterial ligands and nutrients that regulate the host-commensal symbiosis under homeostatic conditions. In addition, *O*-glycosylated mucin from the proximal colon can encapsulate fecal material and the microbiota, sequestering their direct interaction with the mucosa[22]. The Core-1-derived and Core 3-derived complex mucin-type *O*-glycan comprise more than 80% of the murine colon mucin mass [23, 24]. The complex glycans can be further modified through sialyation, fucosylation and sulfation to form terminal epitopes, which is essentially involved in the pathogenesis of IBD)[21]. Proteins such as fucosyltransferase 2 (Fut2) and ST6GALNAC1 (ST6, a dominant sialyltransferase) regulating these terminal glycan epitopes also exert an essential role in maintaining intestinal epithelial homeostasis and microbial symbiosis [25, 26]. Altogether, these data strongly support the premise that both the epithelial layer and the mucus layer are essential in maintaining the intestinal homeostasis, and defects in either increase the susceptibility of developing IBD.

We seek to identify novel pathways and genes that contribute to the pathogenesis of IBD, especially genes that are genetically associated with IBD and preferentially expressed in intestinal epithelial cells. Previously, we and others demonstrated that *C1orf106*, an IBD associated gene that is specifically expressed in intestinal epithelial cells, regulates epithelial cell junctions and permeability [27]. Another group of genes that are genetically associated with IBD and preferentially expressed in the intestinal epithelial cells include nxpe1, nxpe2, and nxpe4 [28]. These genes are located on human chromosome 11 close to the three identified IBD SNPs (rs678170, rs561722, and rs661054), and encoded neuroexphilin and PC-esterase domain (NXPE) family proteins. There are four NXPE family members (NXPE 1-4) in humans. NXPE1 is specifically expressed in colon enterocytes, while the expression of NXPE2 and NXPE4 is detected in both colon and salivary gland [29]. The biological functions of NXPEs are largely unknow.

To understand the potential functions of these protein, we determined the structure of NXPE1 by Cryo-electron microscopy (Cryo-EM). The structure suggests that NXPE1 might act as an *O*-acetyltransferase. Since Nxpe1 is a pseudogene in mice, we further examined this hypothesis by generating a knockout mouse line, in which two homologs of human *NXPE1 gene, Nxpe2* and *Nxpe4* are deleted. Nxpe2 and Nxpe4 share a similar colon expression profile as human NXPE1. We found that mice deficient of Nxpe2 and Nxpe4 completely abolish the *O*-acetylation of sialic acid on the surface of red blood cells, supporting the NXPE family member’s function in regulating *O*-acetylation. These data shed light on the potential pathogenic role of NXPE1 and other family member in IBD.

## Results

### Human and mouse NXPE family members are preferentially expressed in colon

Human *Nxpe1, 2* and *4* are located on chromosome 11, while *Nxpe3* is on chromosome 3. All four human NXPE proteins share similar domain structures consisting of a N-terminal transmembrane domain, followed by a neuroexphilin (NXPH) domain and a C-terminal PC-esterase domain (Figure 1A). NXPE1 is highly homologous to NXPE2 and NXPE4 with sequence identity of 70.1% and 68.89%, respectively. The sequence identity with NXPE3 is significant lower (32.86%). Since several SNPs near human *Nxpe1, Nxpe2*, and *Nxpe4* are associated with UC, we first examined the expression of these genes in various segments of gastrointestinal tract (Figure 1B). The expression of *Nxpe1, 2* and *4* was barely detectible in esophagus, stomach, and different regions of small intestine. Low expression was found in cecum. All three genes are preferentially expressed in various segments of colon, including ascending, transverse, descending and sigmoid colon. Similar expression patterns of these genes are consistent with their genomically clustering in the same locus suggesting a concordant regulation. In contrast, *Nxpe3* from a different chromosome (chr 3) shows high expression only in the esophagus and rectum. Next, we ask whether the expression of *Nxpe1* is impacted by the UC associated SNP rs678170 in the colon [30]. We analyzed the GTEx database with transverse colon samples (Figure 1C). The GG allele resulted in significant lower expression of *Nxpe1* (P value 2.5×10^-20^), which is consistent with the result from a previous publication[30]. As annotated by InterPro [31], The NXPE family proteins contain a putative *N*-terminal transmembrane sequence, followed by a Neurexophilin like domain and a *C*-terminal PC-Esterase domain (Figure 1A). However, whether NXPE1 is an intracellular or an extracellular protein remains unclear. We engineered a FLAG-tag at the *C*-terminal of the NXPE1 and stably expressed it in the HEK293T cell line. FACS analysis with an antibody against FLAG-tag indicated that NXPE1 is expressed on the cell surface, where it could exert its function (Figure 1D). Interestingly, by similar approaches, we only observed lower cell surface expression for NXPE3 and no surface expression of NXPE2 and NXPE4.

**Figure 1.**
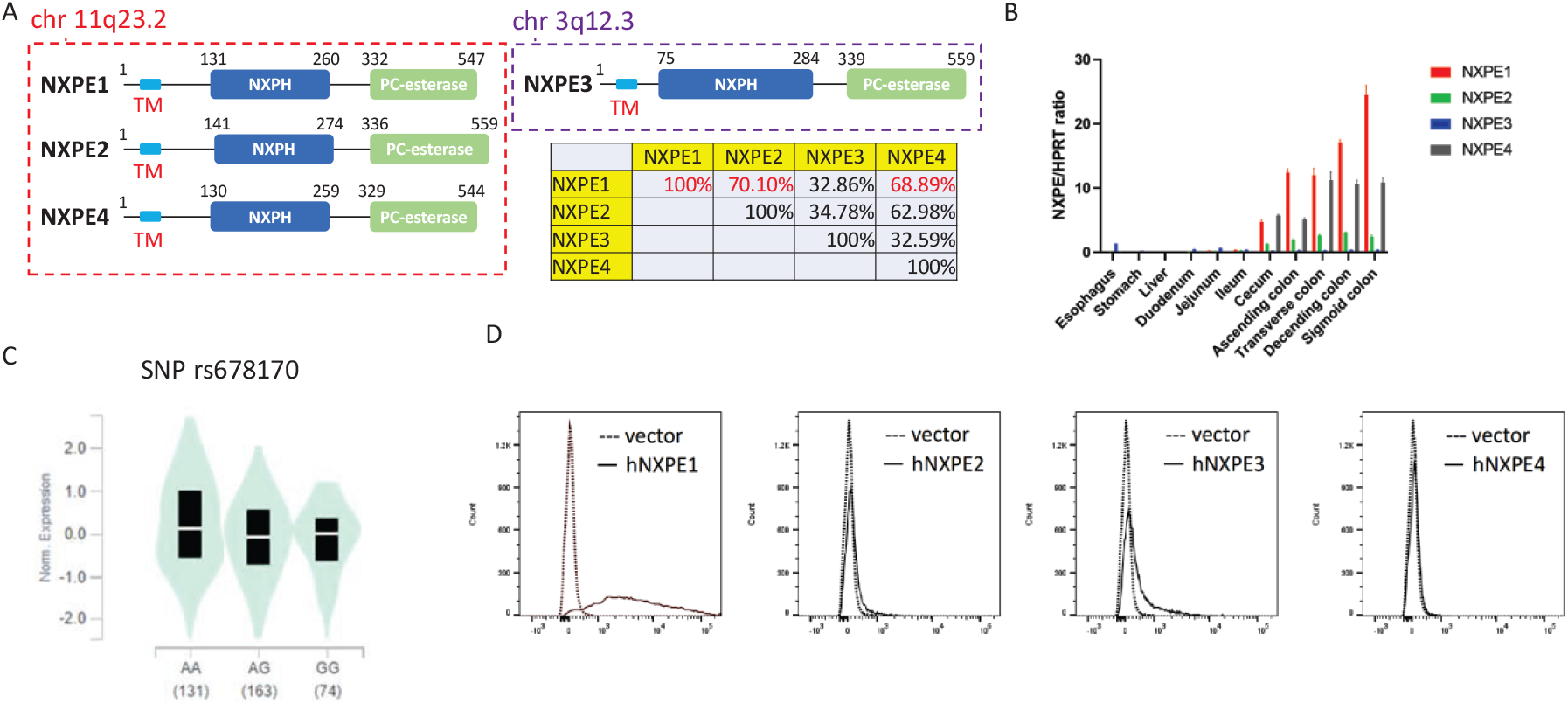
Chromosomal locus and expression location of NXPE family proteins. **A**. Chromosome locus, schematic diagram of domains, and sequence identities of NXPE proteins in human. **B**. The expression of *Nxpe* genes in human gastrointestinal tissues. *Nxpe* expression relative to *Hprt* determined by qPCR is indicated on the y-axis. **C**. Normalized mRNA expression level of *Nxpe1* of SNP rs678170 from GTEx database (P value 2.5×10^-20^). **D**. Cellular colonization of NXPE proteins. HEK293T cells were transfected with expression vectors for human NXPE proteins with C-terminal FLAG-tag. Expression of NXPE proteins were detected by anti-FLAG antibody by FACS.

### Structure of human NXPE1

In order to explore the potential functions of the NXPE family of proteins, we determined the high-resolution structure of the extracellular domain of human NXPE1 (residue 39 – 547) by single particle Cryo-EM at an overall resolution of 2.8 Å (Figure 2, Supplementary Figure 1, Table 1). The high-quality EM map enables accurate model building of a structure containing residues 63 – 547. In the final model, NXPE1 ECD forms a homodimer in C2 symmetry (Figure 2). The cuboid shape has an approximate dimensions of 120 Å X 60 Å X 60 Å. The Loops extending from the N-terminus of both ECD are in proximity and are anticipated to connect to the transmembrane helices (residues 1 – 38).

**Figure 2.**
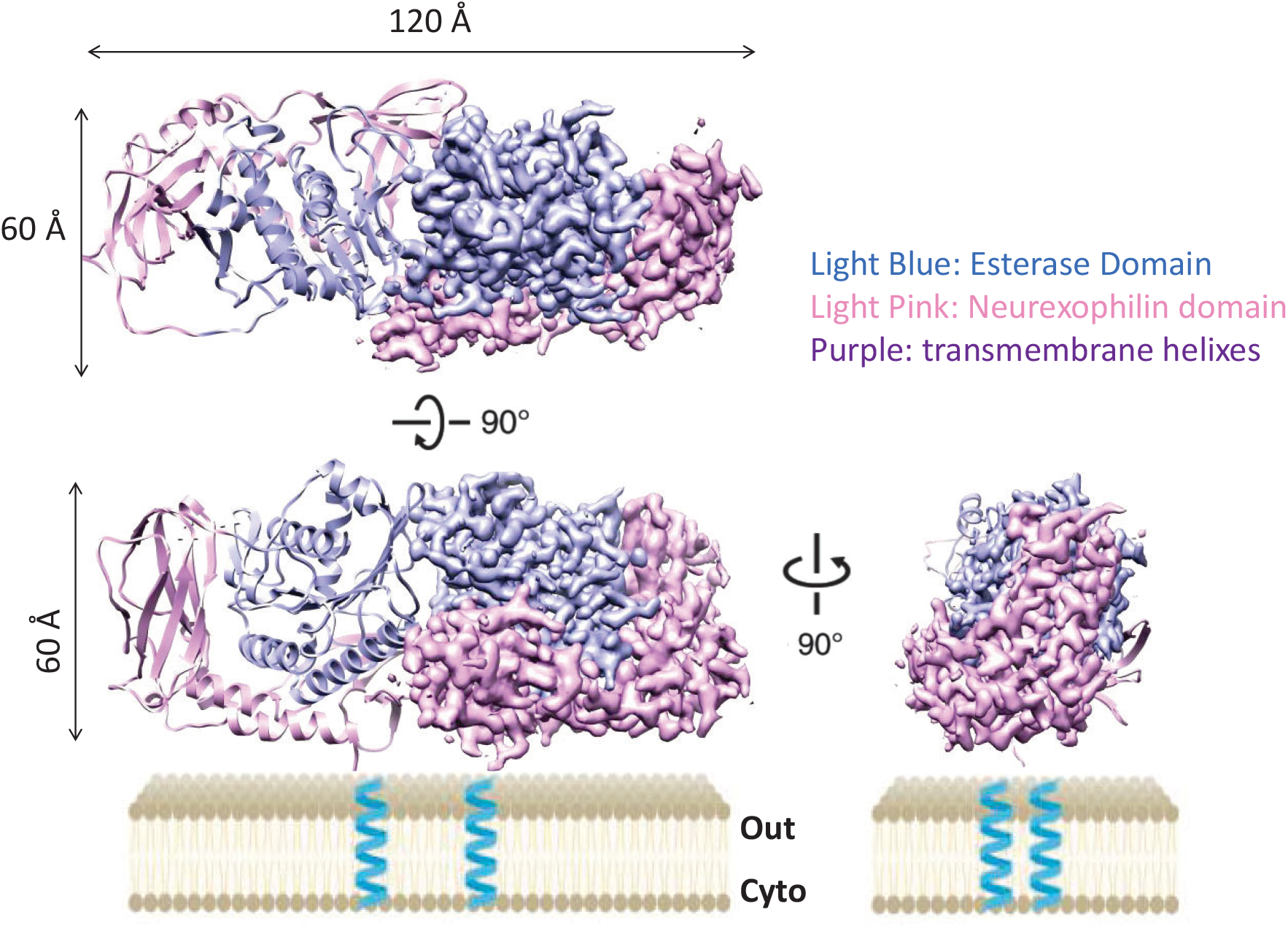
Structure of human NXPE1. viewed from the extracellular side and within the plane of the cell membrane. One protomer is shown in cartoon representation and the other is shown in Cryo-EM density map. The Esterase domain (ED), Neurexophilin domain (ND), and transmembrane helices (shown in the cartoon) are colored in light blue, light pink, and teal (cartoon).

### NXPE1 structure suggests an *O*-acetyltransferase function

NXPE1 exhibits a unique overall fold, with its greatest homology to a published structure (PDB: 6CCI) being only 12% as determined by the Dali server [32, 33]. Each NXPE1 protomer contains an N-terminal Neurexophilin domain (ND) and a C-terminal Esterase domain (ED) (Figure 3A, B). The ED adopts a globular Rossmann fold core, a 7-stranded beta sheet Eβ1 – 7 (red) sandwiched between two sets of helices, set 1 (Eα1,5,6 (orange)) and set2 (Eα2,3,4 (pink)). The ND can be divided into three lobes (ND1 (cyan), ND2 (slate blue), and ND3 (magenta). ND1 and ND3 are both formed by beta sheets and are connected by the ND2 domain consisted of two helices (Figures 3A, B). The three lobes wrap around the esterase domain through contacts around the helices. Despite the tight contacts between the ND and ED domains, a hydrophilic tunnel is formed between ND1 β3,4 from the ND and Eα5,6 from the ED (Figure 3C).

**Figure 3.**
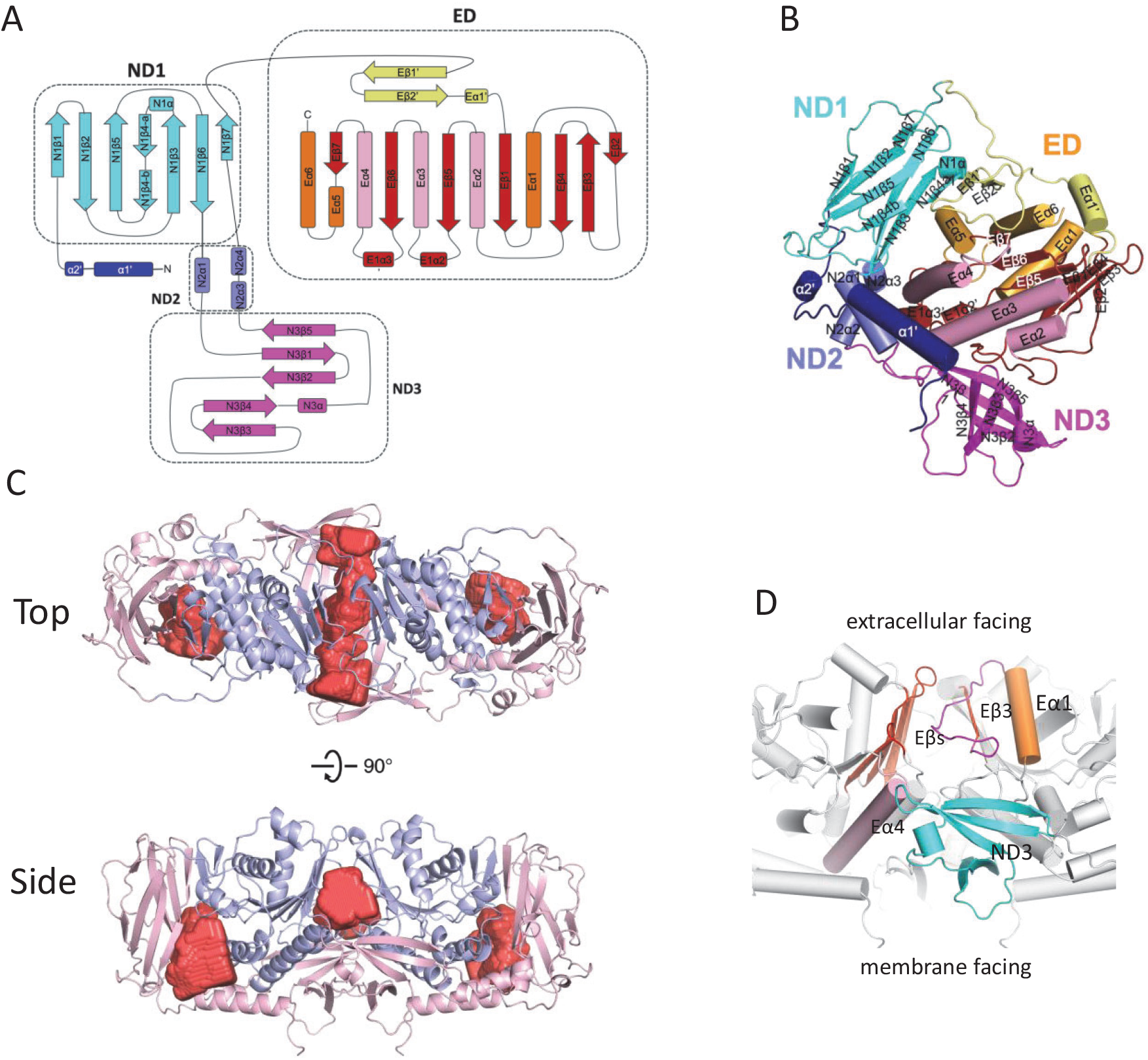
Structural details of human NXPE1. A and B. Topology diagram (A) and monomer structure (B) of NXPE1. The structural elements are color in the same color scheme. C. Hydrophobic cavities shown as surface in the NXPE1 dimers structure, identified by CavityPlus [34]. D. Interactions at the NXPE1 dimer interface.

The two NXPE1 protomers form a homodimer through reciprocal interactions in two regions (Figure 3D). On the extracellular facing side, the long loop formed between the Eα1 and Eβ3 from one protomer closely packs with the edge of the beta-sheet ED from the other protomer (Figure 3D). On the membrane-facing side, the ND3 domains interact with the C-terminus region Eα4 from the opposite protomer (Figure 3D. The contact points carve into a central cavity at the dimer interface (Figure 3C).

We performed a search of secondary structure using PDBeFOLD and identified a distance relative to the esterase domain of NXPE1: XOAT1 (xylan O-acetyltransferase 1, PDB: 6CCI, Figure 4A) [33, 35]. XOAT1 is a xylan *O*-acetyltransferase from *Arabidopsis thaliana* that transfers the acetyl group from acetyl-CoA to xylan. The esterase domain of XOAT1 contains two lobes, a globular lobe that resembles the ED of NXPE1, and a loop-rich lobe. The two lobes form a cleft to allow the binding of substrates such as Acetyl-CoA and xylan. The reaction center is located at the bottom of the cleft, where the Ser-His-Asp triad catalyzes the transferase reaction.

**Figure 4.**
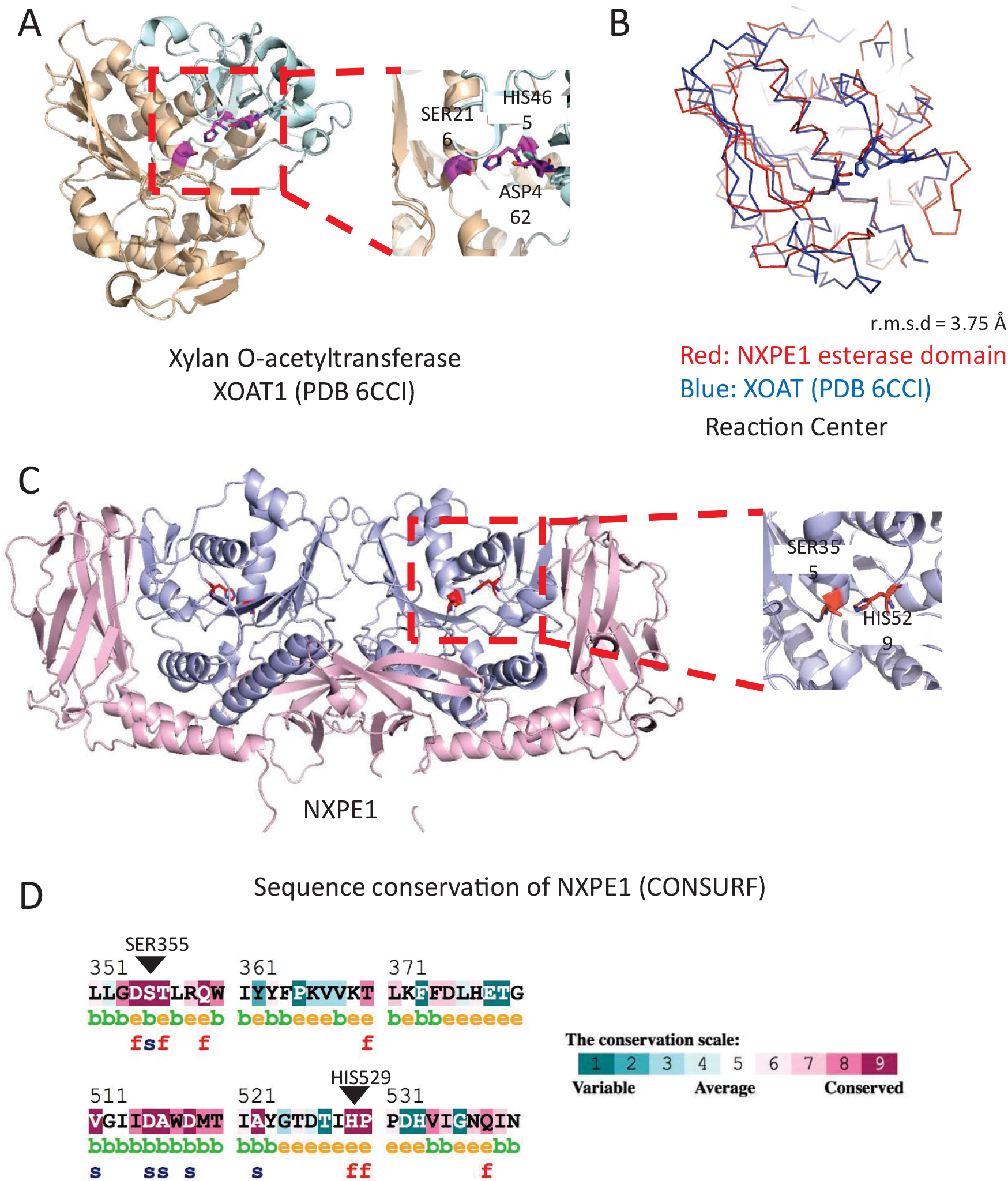
Proposed reaction center in NXPE1. **A**. Structure of XOAT1 (PDB 6CCI) with the globular lobe and loop-rich lobe colored in orange and cyan. The reaction center residues are colored magenta. **B**. Superposition of NXPE1 (red) and XOAT (blue). **C**. Proposed reaction center in NXPE1. **D**. Sequence conservation of NXPE1 calculated by CONSURF.

The Rossman fold of the NXPE1 ED superimposes with XOAT1 with an r.m.s.d of 3.75 Å (Figure 4B). Ser355 and His529 in NXPE1 overlap with Ser216 and His465 from the triad of XOAT1 (Figures 4B and C). These two residues are highly conserved in the NXPE family and throughout the proteins with a similar functional region (CONSURF, Figure 4D) [36]. Hence, we propose Ser355 and His529 are the key residues for sialic acid acetyltransferase activity of NXPE1. We propose reaction mechanism similar to XOAT1. Histidine acts as a general base, abstracting a proton from the serine hydroxyl group and thereby increasing the nucleophilicity of serine. The activated serine then attacks the carbonyl carbon of acetyl-CoA, resulting in the formation of a tetrahedral intermediate and a covalent acetyl-serine intermediate, with the release of CoA. Next, sialic acid, as the acceptor molecule, attacks this acetyl-enzyme intermediate. Histidine again functions as a general base to facilitate the transfer, resulting in the acetyl group being transferred from serine to sialic acid. However, different from XOAT1, there is no aspartate residue adjacent to His529 in NXPE1. In XOAT1, the aspartate residue forms hydrogen bond to stabilize and orient histidine. Aspartate is less conserved in the reaction center, as Ser-His dyad has also been reported.

Capsule structure 1 domain 1 (CASD1) functions as a sialate O-acetyltransferase, a crucial enzyme involved in the biosynthesis of 9-O-acetylated sialoglycans. It catalyzes the 9-O-acetylation of terminal sialic acid sugars on glycoproteins and glycolipids. [37]. The esterase domain of NXPE1 has 80.2 % sequence identity to that of CASD1 (Supplementary Figure 2). We thus compared the structure of NXPE1 with the AlphaFold model of the esterase domain of CASD1 (Supplementary Figure 4A). All secondary structures of CASD1 are well superimposed with NXPE1 (r.m.s.d = 3.0) (Supplementary Figure 4). In the reaction center, Ser355 and His529 from NXPE1 superimposed with Ser94 and His273 from CASD1, whereas ASP270 from CASD1 is also in close proximity, form the Ser-His-Asp triad like the one from XOAT1 [37].

The ND in NXPE1 assumes a unique structural arrangement. In the crystal structure of Neutexopilin-1 in complex with neurexin-1 LNS2 domain (6PNQ), in which Neutexopilin-1 forms a single domain with two beta-sheets packing against each other (Supplementary Figure 4B) [38]. Whereas, in NXPE1, the insertion of two helices segregates the Neutexopilin domain into 3 lobes. Whether the neurexophilin domain has the potential to adopt different conformations is to be studied. These data suggest that the Neurexophilin domain may assume different conformations to regulate the *O*-acetyltransferase functions or mediate additional functions such as protein-protein interactions that regulate the function of NXPE1.

### Double knockout of Mouse NXPE2 and NXPE4 results in lack of sialic acid *O*-acetyltransferases function

Given the sequence similarity between NXPE1 and CASD1 EDs, we hypothesized that these two proteins may have common biological functions. The *Nxpe1* gene in mouse is a pseudogene. However, human and mouse *Nxpe2* and *Nxpe4* genes share very high sequence homology with the human *Nxpe1* gene (Figure 5A, Supplementary figure 3). We speculated that mouse NXPE2 and NXPE4 proteins might have similar functions as the human NXPE1. We therefore first examined the expression of mouse *Nxpe2* and *Nxpe4* in the gastrointestinal tract (Figure 5B). Similar to that of human *Nxpe1*, mouse *Nxpe2* and *Nxpe4* were not detected in esophagus, stomach and small intestine. In contrast to human *Nxpe1*, the expression of *mNxpe2* and 4 were not detected in cecum. Low expression of *mNxpe2*, but not *mNxpe4*, was observed in proximal colon. Both *mNxpe2* and *4* were highly expressed in middle and distal colon. The expression of *mNxpe2* is also detected in liver. In addition, when tagged *mNXPE2* and *mNXPE4* were expressed in HEK293T cells, they could be stained on cell surface (Figure 5C) similar to hNXPE1. All these data together supported that mNXPE2 and mNXPE4 could be used to explore the functions of the NXPE family proteins.

**Figure 5.**
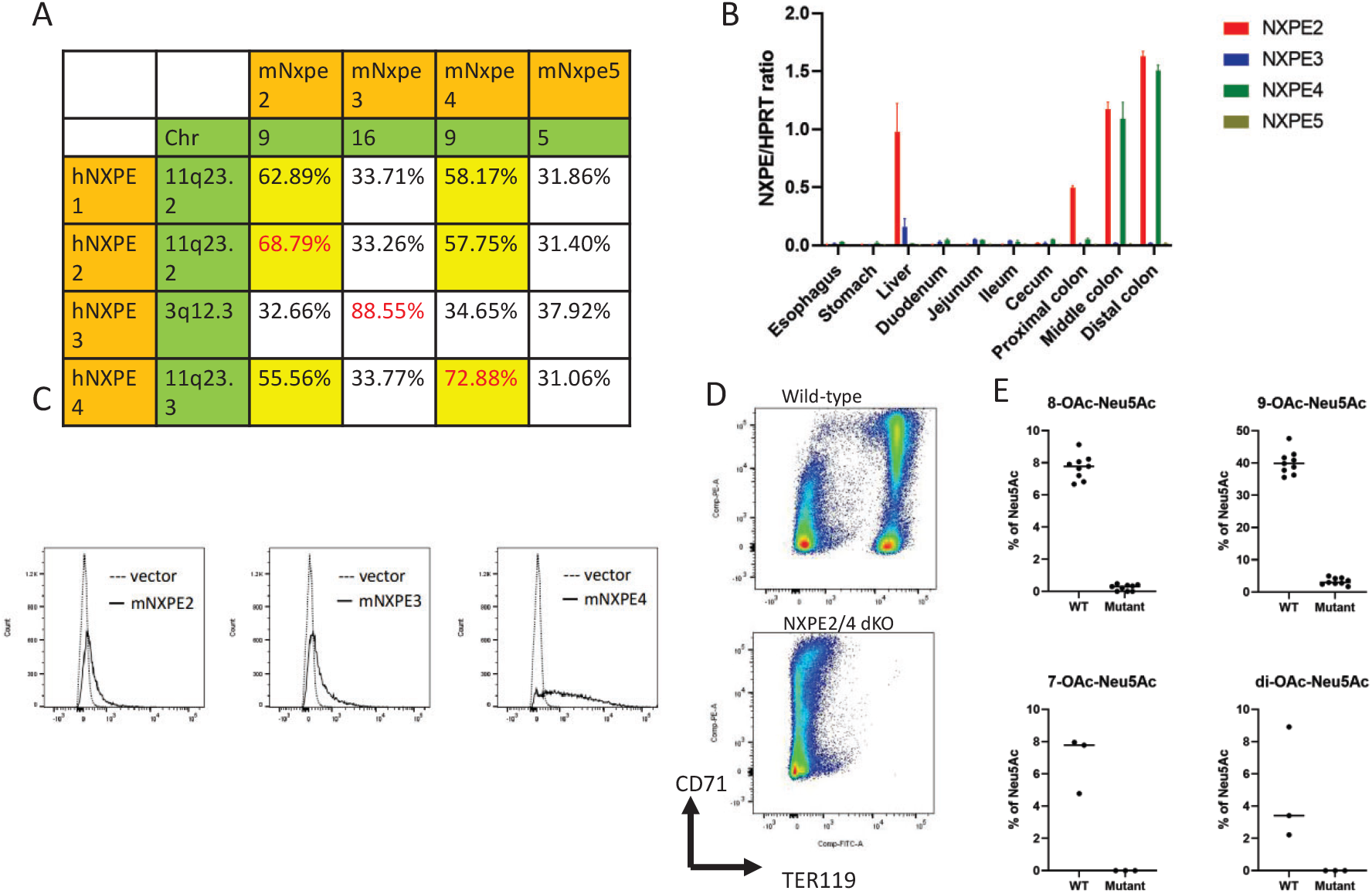
Nxpe2/Nxpe4 dKO mouse and phenotypical characterizations. **A**. Protein sequence identities between human and mouse NXPE proteins. **B**. The expression of mouse Nxpe2, Nxpe3, Nxpe4, and Nxpe5, in gastrointestinal tissues. The relative expression was plotted as the ratio of Nxpe/Hprt. Total RNA was extracted from C57Bl/6 mice. NXPE2, NXPE3, NXPE4 and NXPE5 qPCR were done in QuantStudio 7 Flex. **C**. Cellular colonization of NXPE. HEK293T cells were transfected with expression vectors for mouse proteins with C-terminal FLAG tag. Expression of NXPE proteins were detected by anti-FLAG antibody by FACS. **D**. TER119 marker profiling of RBC cells. The RBC are isolated, and cell surface makers CD71 and TER119 are detected by FACS. **E**. O_Ac_-Neu5Ac glycan species profiling to the extracted RBC membranes by UPLC.

Since mNXPE2 and mNXPE4 share high sequence identity and similar expression patterns, they may perform redundant functions. We generated double knockout mouse line in which both *mNxpe2* and *mNxpe4* genes are deleted (dKO NXPE2/4, Figure 5D) to evaluate the potential roles of both proteins together. These dKO mice developed normally and with no obvious defect in the gastrointestinal tract and immune system. Previous studies have demonstrated that CASD1 is a sialic acid 9-O-acetyl transferase[37]. The red blood cells (RBCs) from CASD1-/-mice lose their ability to be bound by a specific lineage detecting antibody, TER-119 [39]. The epitopes detected by TER-119 has been suggested to be a 7,9-di-O-acetyle form of sialic acid on RBC surface [39]. We therefore examined whether NXPE2/4 dKO impacted the TER-119 epitopes on the RBCs. Interestingly, the binding of TER-119 to RBCs and their progenitors was completely abolished, supporting that either mNXPE2, 4 or both also participate in the O-acetylation of sialic acid in RBCs as does CASD1 (Figure 5D).

The acetylation of salic acid can happen at position 4, 7, 8, and 9. To further investigate the impacts of NXPE2/4 dKO on the acetylation of salic acid on RBCs, we used UPLC to compare the difference of the cell surface glycan profile between purified RBC cell membranes of NXPE2/4 dKO and wild-type mice. As shown in Figure 5E and Supplementary Figure 5, in WT mice, 9’ O-acetylated sialic acid is the major form detected, followed by 8’ and 7’- O-acetylated sialic acid. Low amounts of di-acetylated sialic acid was detected in WT mice. However, all these forms were drastically reduced in dKO NXPE2/4 mice, confirming an indispensable role of mNXPE2/4 in the O-acetylation of sialic acid at these positions in RBCs. Together, these data support that NXPE family proteins possess the functions of regulating sialic acid O-acetylation.

## Discussion

To summarize, we determined the high-resolution structure of human NXPE1 by Cryo-EM. Based on the analysis of structural similarity to XOAT, we propose that NXPE1 is an *O*-acetyltransferase with the reaction center at Ser355 and His529. However, the reaction center is not adjacent to a substrate molecule binding pocket. Hence, potential conformational changes or protein binding partners that bring the substrate to the reaction center are to be studied. Interestingly, the ND of NXPE1 does not form a single beta-sheet bundle as observed in the XOAT homolog structure determined previously. Instead, the ND segregates into 3 lobes and wraps around the ED. Potential conformational changes and the effects of these changes on the esterase activity are to be studied.

Overall, this structure provides valuable insights into the reaction mechanism of NXPE1’s O-acetyltransferase activity. Based on its similarity to previously characterized enzymes such as XOAT1, we propose that NXPE1 contains a catalytic center composed of a His529-Ser355 dyad, where histidine acts as a general base and serine functions as a nucleophile. Unlike the classical His-Ser-Asp triad, NXPE1 lacks an aspartate residue to stabilize and orient histidine. However, similar dyad catalytic centers have been reported in other enzymes, such as serotonin N-acetyltransferase, rhomboid protease, and nucleoporin [40, 41]. The mechanism by which histidine is stabilized in NXPE1 remains to be investigated.

Sialic acid is normally the terminating units of glycan chains in glycoproteins and consists of a family of nine carbon sugars, in which 5-*N*-acetyl-neuraminic acid (Neu5Ac) is the most common form in human [42]. As we discussed previously, studies have established a critical role of sialylation of terminal glycans in maintaining the integrity of the mucus layer and preventing the development of IBD[25, 26]. Sialic acid Neu5Ac can be further modified to form diverse family members, with *O*-acetylation widely observed[43]. *O*-acetylation happens at the C-4, 7, 8, 9 position of Neu5AC, and sialic acid-specific O-acetyltransferases (SOATs) and O-acetylesterases (SIAEs) that add and remove O-acetyl groups, respectively. CASD1, thus far, is the only human enzyme identified to catalyze the O-acetylation of Neu5Ac at the position C-7 [37, 44]. A hypothetical migrate has been proposed to migrate the acetyl group from C-7 to C-8 and C-9 positions, but the enzyme has yet to be identified. Ter-119 is a rat antibody that recognize the 7,9-di-O-acetyl form of sialic acid glycoform on erythrocyte lineage. CASD1 knockout mice completely abolished the recognition of epitope on surface of red blood cell.

In our current study, we observed similar phenotypes in a mouse NXPE2 and NXPE4 double knockout. In addition, by profiling the terminal glycan profile, we found that the O-acetylation species on Neu5Ac including, 7-OAc-Neu5Ac, 8-OAc-Neu5Ac, 9-OAc-Neu5Ac and di-OAc-Neu5Ac, all drastically reduced to level of baseline in the double knockout mice (Figure 5E), supporting an indispensable role of NXPE2 and NXPE4 in the O-acetylation of sialic acid. Mouse NXPE2 and NXPE4 proteins share high sequence homology (Supplementary Figure 3) and both are expressed in the colon, so it is difficult to identify which protein or if both contribute to the O-acetylation process. A recent study, however, using mNxpe2 or mNxpe4 overexpressing cell lines, identified Nxpe2 overexpression but not that of Nxpe4 conferred the Ter119 binding to Glycophorin A[45], suggesting that Nxpe2 knockout contributes to the phenotype describe in this study.

Human NXPE1 is a pseudo gene in mouse, but its sequence is also homologous to both mouse Nxpe2 and Nxpe4. Based on the structure data and mouse knockout data, we believe NXPE1 also processes O-acetyltransferase function in human. This hypothesis is supported by two recent studies posted online that human NXPE1 but not NXPE4 catalyzed O-acetylation of Neu5Ac on Mucin protein[46, 47]. NXPE1 specifically modifies the 9-OH group to produce 9-O-acetylated Neu5Acm, and is capable of transferring the acetyl group from acetyl-coenzyme A to sialic acid *in vitro*. Interestingly, NXPE1 has been identified as the key enzyme to regulate the mild periodic acid Schiff staining (mPAS) on colon tissues through the modification of acetylation of sialic acid[47]. Importantly, a protective mutation (G353R) in humans drastically reduced the acetylation of Neu5Ac on mucins. In a study evaluating expression quantitative trait loci (eQTL) in the human colon, the expression of NXPE1 is identified as one of the target genes of UC-associated SNP (rs678170). The protective GG genotype results in significantly lower expression of NXPE1 in the colon than that from AA genotype [30]. Altogether, these data suggest the expression and function of NXPE1 impacts the pathogenesis of UC through the regulation of acetylation of salic acid. Our detailed structural data provide essential information for developing novel therapies for the treatment of IBD.

## Methods

### 1. Expression analyses of *Nxpe* genes in gastrointestinal tissues

RNA samples of human gastrointestinal tissues were obtained from Amsbio Inc., and mouse total RNA were extracted from C57Bl/6 mice. The expression of human Nxpe1, Nxpe2, Nxpe3, Nxpe4, mouse Nxpe2, Nxpe3, Nxpe4, and Nxpe5 were analyzed by qPCR in QuantStudio 7 Flex. *Hprt* was used as a control. The relative expression was plot as the ratio of Nxpe/Hprt. The normalized mRNA expression of Nxpe1 associated with different alleles of SNP rs678170 was obtained by analyzing the GTEx database.

### 2. Nxpe2/Nxpe4 dKO mouse generation

The *Nxpe2 & Nxpe4 double* knockout mouse were generated and provided by Biocytogen. In brief, two sgRNAs were designed to target a region upstream of *Nxpe2* gene exon3 and downstream of *Nxpe4* gene. Cas9 mRNA and sgRNAs were mixed and co-injected into the cytoplasm of one-cell stage fertilized eggs obtained from C57BL/6N female mice. After injection, surviving zygotes were transferred into oviducts of KM albino pseudopregnant females to generate F0 founders. F0 founder mice with expected genotype were confirmed by tail genomic DNA PCR and sequencing and were mated with C57BL/6N mice to generate germline-transmission confirmed F1 heterozygous mice. The completely knock out of both Nxpe2 and Nxpe4 was confirmed by the absence of the mRNA expression of Nxpe2 and Nxpe4 in the knockout colon tissue (data not shown).

gRNA sequence:

GAATACATTATAGGCAGTAC**TGG**

AGTTCTTGTAAGGCGGGCCA**AGG**

### 3. Protein cell surface expression analyses by Flow cytometry

For sub-cellular expression location determination, NXPE genes in pTT5 vectors are transfected to HEK293T cells. After 2 days single cell suspensions were prepared for flow cytometry analysis using Anti-FLAG PE antibody.

For TER119 marking profiling to mouse RBC, mouse blood was centrifugated at 1000 g for 5 minutes to form gradient layers by cell types, and the red blood cell layers are isolated, and washed with PBS buffer. Single cell suspensions were prepared and flow cryometry analysis was performed using TER119 FITC antibody

### 4. Cell membrane glycan profiling

Blood was collected from mouse. The blood was centrifugated at 1000 g for 5 minutes to form gradient layers by cell types, and the red blood cell layers are isolated, and washed with PBS buffer. To isolate the cell membrane, the red blood cells was first treated by hypotonic buffer (5 mM NaCl, 20 mM HEPEs pH 7.4) for cell lysis. After 10,000 g centrifugation, the pellet was isolated and washed by hypertonic buffer (500 mM NaCl, 20 mM HEPEs pH 7.4). After another centrifugation, the membrane pellet was resuspended in dH2O.

Samples were suspended in ultrapure water (Invitrogen, USA) containing 1µl of protease inhibitor (Protease Inhibitor Cocktail Set III, EDTA-Free, Cat# 539134, EMD Millipore Corp, Germany) and then sonicated for two pulses of 15s each. Protein estimation was done using the Pierce BCA Protein assay Kit (Thermo Scientific, Rockford, IL, USA). Known amount of sample was hydrolyzed with 2M glacial acetic acid (Sigma) at 80°C for 3h followed by removal of excess acid using speed vacuum. DMB (1,5-diamino-4,5 methylenedioxybenzene, dihydrochloride, Mol wt. 225.07, Code M021, Dojindo Lab) derivatization was performed as reported previously (1,2,3) and analysis was done using RP-UPLC-FL (Waters, Acquity UPLC) system on BEH C18 column (2.1 × 50 mm x 1.7µm, Waters). Sialic acids (Neu5Ac, Neu5Gc and O-acetylated forms) were eluted with two solvent mixtures of Solvent-A (7% methanol HPLC grade, Sigma containing 0.1% TFA), in water and Solvent-B (100% acetonitrile HPLC grade, Sigma containing 0.1% TFA) at a flow rate of 0.4 mL/min. The excitation and emission wavelengths were 373nm and 448nm, respectively. Known amount of standards were used to quantify the sialic acids present in the samples[48-50].

### 5. NXPE construct design and vector cloning

For sub-cellular expression location determination, full length NXPE genes from human and mouse was cloned into pTT5 vectors, fused with c-terminal FLAG tag. For structure determination, the DNA sequence of the secretion signal MDMRVPAQLLGLLLLWLPGARC followed by a FLAG tag and a TEV cleavage site was fused to the N-terminal of residue 39 – 547 of human NXPE1, and cloned into pTT5 vectors.

### 6. Expression and purification of human NXPE1

NXPE1 (39-547) vector was transfected to expi293 GnTI^-^ cells. The cells were shaken in 8% CO_2_, 80% humidity, and 37 °C for 5 days. The cell supernatant was harvested by removal of the cell pellet by 1500 g centrifugation. To purify the protein, the cell supernatant was incubated with FLAG M2 agarose beads for 2 hours at 4 °C. The beads were collected by centrifugation, washed by buffer A (150 mM NaCl, 20 mM HEPEs pH 7.4), and eluted by buffer A plus 150 ng/µL Flag peptides. The protein was further purified by size exclusion chromatography with a Superdex 75 column, in buffer A.

### 7. Cryo-EM sample preparation and data acquisition

Because of a strong preferred orientation issue, the cryo-EM grids were prepared in two ways. First, the Chameleon system was used. SPT Labtech 300 mesh Cu R 1.2/1.3 holey carbon self-wicking nanowire grids were glow discharged with a Pelco easiGlow device using mixed air at 15 mA for 45 seconds. 0.1 mg/ml NXPE1 sample was applied to the grid and vitrified by Chameleon at 100 ms. These grids are used to collect the untitled and 30% tilt data sets. Second, the detergent octyl glucoside (OG) was added to the final sample before freezing to further correct the preferred orientation. Previously frozen NXPE1 protein was diluted to 3 mg/ml while 0.1 % OG was added to the final sample. 2.5 uL of sample was then applied to Quantifoil® holey carbon 1.2/1.3-Au 300 mesh grids glow-discharged with a Pelco easiGlow device using mixed air at 15 mA for 45 seconds. Grids were blotted with grade 595 standard Vitrobot filter paper for 5 s at 4 °C and 100% humidity using a Vitrobot® Mark IV, followed by rapid plunging into liquid ethane cooled by liquid nitrogen. These grids are used to collect the 0.1% OG data set. Dose fractionated movies with a pixel size of 0.813 Å per pixel were recorded using EPU software at defocus ranging from -1.0 to -2.0 μm with a total electron exposure of 54 e^−^/ Å^2^.

### 8. Image processing

CryoSPARC is used for all data processing steps[51]. Micrographs were corrected for beam-induced drift using Patchmotion [52]. The contrast transfer function (CTF) parameters for each micrograph were determined using PatchCTF [52]. Details on the data processing work flow are shown in Supplementary figure 1. Briefly, 3 datasets (untilted, 30° tilt, and 0.1 % OG) was combined for the final structure. After iterative 2D and 3D classification and refinement, a total of 106,947 particles were retained in the final dataset, which reached a nominal resolution of 2.82 Å. Statistics of the final map are calculated using the Fourier shell correlation (FSC) criterion and a threshold of 0.143 using phenix_comprehensive_validation tool and reported in Table S1.

### 9. Model building and structure refinement

AlphaFold model of the NXPE1 monomer (AF-Q8N323-F1-model_v1) was used as the initial model after removing the N-terminal helix [53]. The structure was then refined in Coot [54] and in phenix.real_space_refine [55]. Statistics of the model are reported in Table S1. Figures were prepared with Pymol (The PyMOL Molecular Graphics System, Version 2.0 Schrödinger, LLC) and chimera [56].

## Supplementary Information

**Table S1.** Cryo-EM data collection, refinement and validation statistics.

| NXPE1/(EMDB-XXXXX)/(PDB XXXX) |  |
| --- | --- |
| **Data collection and processing** |  |
| Microscope | FEI Titan Krios |
| Camera | Gatan K3 direct detector |
| Magnification | 105000 |
| GIF slit width (eV) | 20 |
| Voltage (kV) | 300 |
| Electron exposure (e<sup>-</sup>/Å<sup>2</sup>) | ~58 |
| Exposure rate (e<sup>-</sup>/pixel/s) | ~24 |
| Number of frames per micrograph | 50 |
| Automation software | EPU |
| Defocus range (μm) | 1.0 - 2.0 |
| Pixel size (Å) | 0.813 |
| Symmetry | C1 |
| Number of movies | 9149 |
| Initial particle images (no.) | 14,158,967 |
| Refined particle images (no.) | 7,001,742 |
| Final particle images (no.) | 106,947 |
| Map resolution (Å) (masked/unmasked) | 2.8/3.0 |
| FSC threshold | 0.143 |
| Map resolution range (Å) | 2.0 – 3.0 |
| **Refinement** |  |
| Refinement package | Phenix-1.19.2-4158 |
| Initial model used (PDB code) | AlphaFold2 |
| Model resolution (Å) | 3.2 |
| FSC threshold | 0.5 |
| Map sharpening *B* factor (Å<sup>2</sup>) | -75.6 |
| Model vs map CC | 0.83 |
| Model composition |  |
| Non-hydrogen atoms | 8077 |
| Protein residues | 970 |
| Ligands | 2 |
| *B* factors (Å<sup>2</sup>) |  |
| Protein | 114.9 |
| Ligand | 91.16 |
| R.m.s. deviations |  |
| Bond lengths (Å) | 0.003 (0) |
| Bond angles (°) | 0.555 (0) |
| Validation |  |
| MolProbity score | 1.70 |
| Clashscore | 10.05 |
| Poor rotamers (%) | 0.23 |
| CaBLAM outliers (%) | 1.14 |
| Cβ deviation | 0 |
| Ramachandran plot |  |
| Favored (%) | 97.00 |
| Allowed (%) | 3.00 |
| Disallowed (%) | 0.00 |

