## Supplementary Figures for "Structure and function of human NXPE1, a sialic acid *O*-acetyltransferase"

**Supplementary Figure 1. Cryo-EM data processing workflow**

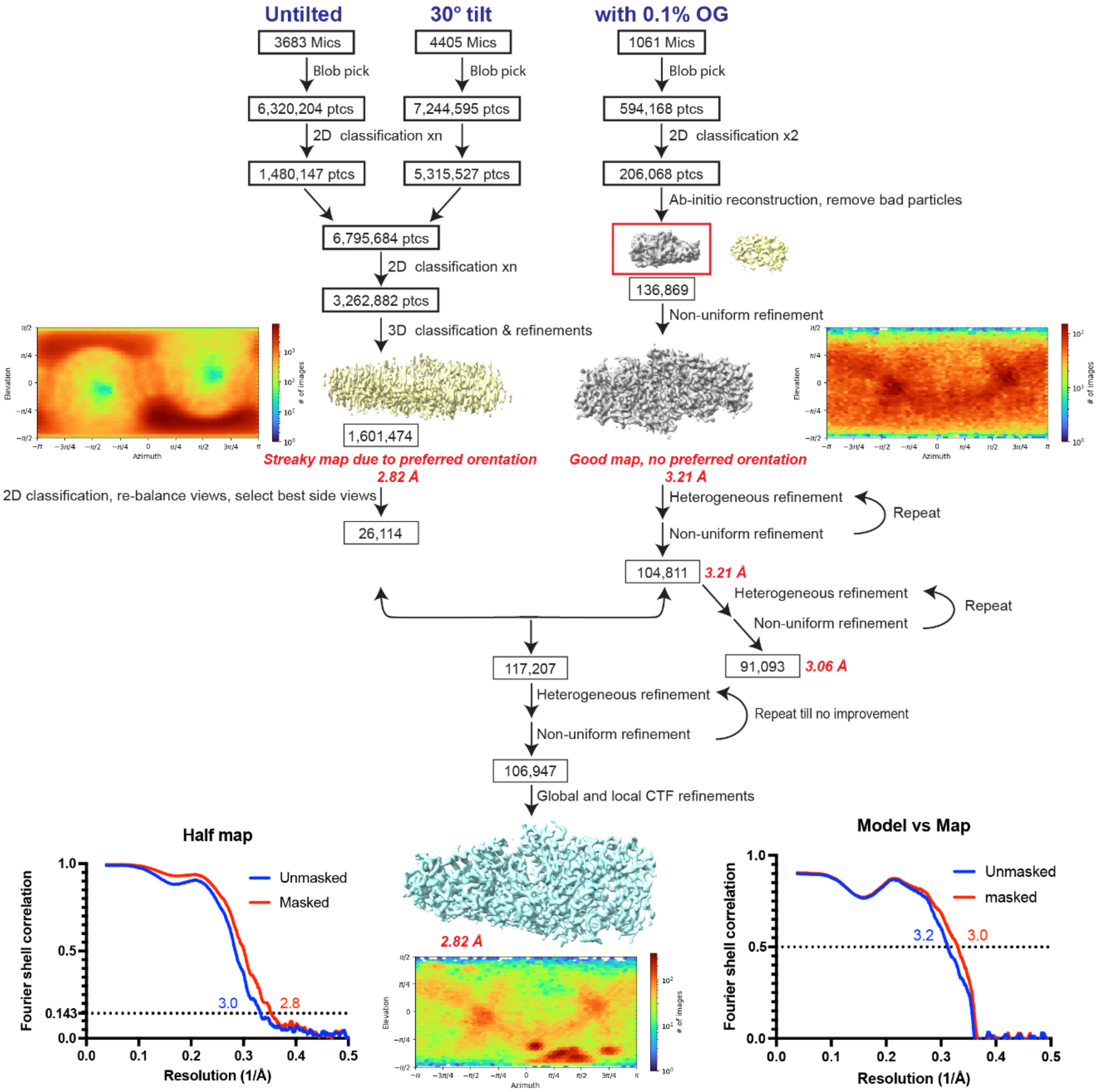

**Supplementary Figure 2. Sequence alignment of human NXPE1 and CASD1**

```
CASD1       NDSCEYLL---SSGRFLGEKVWQPHSCMMHKYKISEAKNCLVDKHIAFIGDSRIRQLFYS     57
NXPE1       EETCQVGMKPPVPGGYTLQGKWITTFCNQVQLDTIKINGCLKGKLIYLLGDSTLRQWIYY     60
            :::*:  :     * :  :  *    *   : .  : :.** .* * ::*** :** :*

CASD1       FVKIINPQF----KEEGNKHENIPFEDKTASVKVDFLWHPEVNGSMK-QCIKVWTEDSIA     112
NXPE1       FPKVVKTLKFFDLHETGIFKKHLLLDA---ERHTQIQWKKHSYPFVTFQLYSLIDHDYIP     117
            * *:::       :* *  :::: ::    . :.:: *: .    :. *  .:  .* *

CASD1       KPHVIVAGAATWSIKIHNGSSEALSQYKMNIT-----SIAPLLEKLAKTSDVYWVLQDPV     167
NXPE1       REIDRLSGDKNTAIVITFGQH--FRPFPIDIFIRRAIGVQKAIERL--------FLRSPA     167
            :    ::*  . :* *  *.   :  : ::*      .:   :*:*        .*:.*.

CASD1       YEDLLS-EN-RK-MITNEKIDAYNEAAVSILNSSTRNSKSNVKMFSVSKLIAQETIMESL     224
NXPE1       TKVIIKTENIREMHIETERFGDFHGYIHYLIMK-DIFKDLNVGIIDA----WDMTIAYGT     222
             : ::. ** *:  * .*::. ::     :: .    .. ** ::..     : **  .

CASD1       DGLHLPESSRETTAMILMNVYCNKILKPVDGSCCQP 260
NXPE1       DTIHPPDHVIGNQINMFLNYIC-------------- 244
            * :* *:    .   :::*  *
```

**Supplementary Figure 3. Sequence alignment of mouse and human NXPEs**

```
Mouse_NXPE3      ------------------MQ-LTRCCFVFLVQGSL-------------------------         16
Human_NXPE3      -------------MWTNFFKLRLFCCLLAVLMV-VVLV--INVTQVEYLDHETVSATFID         44
Human_NXPE4      -------------MKISMINYKSLLALLFILASWIIFTVFQNSTKVWSALNLSISLHYWN         47
Mouse_NXPE4      -------------MKISMINYKSLLALLFILASWIIFTVFQNSTKVWSALNLSISLHYWN         47
Mouse_NXPE2      -MRRMLSPRILLSSLPNASARKLFLIVLIIFVFWVVFMTSKDHTEFMVHLNNRIILRRWS         59
Human_NXPE1      ------------MSSNTMLQKTLLILISFSVVTWMIFIISQNFTKLWSALNLSISVHYWN         48
Human_NXPE2      MVEKILI-HRILTLFPNAIARKLLLMLTFILIFWIIYLASKDHTKFSFNLENHIILNQGN         59
                                           .   .   :

Mouse_NXPE3      --------------YLVICGQDDGPPGSED-PEHDDHE---GQPRPR-VPRKRGHISPKS         57
Human_NXPE3      SSGQFVSSQVTGISRNPYCGYDQQTLSSQERMEEDSLLAALHRQVPDVGPVPFVK---ST         101
Human_NXPE4      NSTKSL------FPKTPLISLKP---LTETELRIKEIIEKLDQQIP---PRPFTHVNTTT         95
Mouse_NXPE4      NSTKSL------FPKTPLISLKP---LTETELRIKEIIEKLDQQIP---PRPFTHVNTTT         95
Mouse_NXPE2      IFKEFL------HSEELK-NTPA---SVEAELAVTAILEKLNQQIP---PRPFQTHSSTT         106
Human_NXPE1      NSAKSL------FPKTSLIPLKP---LTETELRIKEIIEKLDQQIP---PRPFTHVNTTT         96
Human_NXPE2      IFKKYS------HSETPLCPAVS---PKETELRIKDIMEKLDQQIP---PRPFTHVNTTT         107
                                             :             :  *   *        .:

Mouse_NXPE3      RPLANSTLLGLLAPPGEVWGVLGQPPNRPKQSPLPSTKVKKIFGWGDFYSNIKTVALNLL         117
Human_NXPE3      DP--SSSYFVILN-SAAFFKVGSQLEVLV--HVQDFQRKPKKYGGDYLQARIHSLKLQAG         156
Human_NXPE4      SA--THSTATILN-PRDTYCRGDQLHILL--EVRDHLGRRKQYGGDFLRARMSSPALMAG         150
Mouse_NXPE4      SA--THSTATILN-PRDTYCRGDQLHILL--EVRDHLGRRKQYGGDFLRARMSSPALMAG         150
Mouse_NXPE2      SA--KQSTATIHN-PQRTYCVGDQLNVLL--VAKDYFGNRKEYGGDFLRARIFSPAMKAG         161
Human_NXPE1      SA--THSTATILN-PRDTYCRGDQLDILL--EVRDHLGQRKQYGGDFLRARMSSPALTAG         151
Human_NXPE2      SA--THSTATILN-PQDTYCRGDQLDILL--EVRDHLGHRKQYGGDFLRARMYSTALMAG         162
                     . :   :       :   .*                * :* . : :.: :  :

Mouse_NXPE3      VTGKIVDHGNGTFSVHFRHNATGQGNISISLVPPSKAVEFHQEQQIFIEAKASKIFNCRM         177
Human_NXPE3      AVGRVVDYQNGFYKVFFTLLWPGKVKVSVSLVHPSEGIRVLQRLQE--DKPDRVYFKSLF         214
Human_NXPE4      ASGKVTDFNNGTYLVSFTLFWEGQVSLSLLLIHPSEGVSALWSARN--QGYDRVIFTGQF         208
Mouse_NXPE4      ASGKVTDFNNGTYLVSFTLFWEGQVSLSLLLIHPSEGVSALWSARN--QGYDRVIFTGQF         208
Mouse_NXPE2      TSGKVTDFNNGTYLVSFTLFWEGPVSLSILLMHPSEGVSALWRARN--RGYGKIIFTGQF         219
Human_NXPE1      ASGKVMDFNNGTYLVSFTLFWEGQVSLSLLLIHPSEGASALWRARN--QGYDKIIFKGKF         209
Human_NXPE2      ASGKVTDFNNGTYLVSFTLFWEGQVSLSLLLIHPSEGVSALWRARN--QGCDRIIFTGLF         220
                 . *:: *. ** : * *     *  .:*: *: **:.       :          *.  :

Mouse_NXPE3      EWEKVERGRRTSLCTHDPAKICSRDHAQSSATWSCSQPFKVVCVYIAFYSTDYRLVQKVC         237
Human_NXPE3      RSGRISETTECNVCLPGNLPLCNFTDLYTGEPWFCFKPKKLPCSSRITHFKGGYLKGLLT         274
Human_NXPE4      VNGTSQVHSECGLILNTNAELCQYLDNRDQEGFYCVRPQHMPCAALTHMYSKNKKVSYLS         268
Mouse_NXPE4      VNGTSQVHSECGLILNTNAELCQYLDNRDQEGFYCVRPQHMPCAALTHMYSKNKKVSYLS         268
Mouse_NXPE2      LNGTSPVLTECGLTLNTSAELCQYLDARDHEAFYCLKLPGVPCEALTHMTSKNSNISYLS         279
Human_NXPE1      VNGTSHVFTECGLTLNSNAELCEYLDDRDQEAFYCMKPQHMPCEALTYMTTRNREVSYLT         269
Human_NXPE2      ANRSSNVFTECGLTLNTNAELCQYMDDRDQEAFYCVRPQHMPCEALTHMTTRTRNISYLS         280
                          . .:       :*.  .      : * :   : *       .       :

Mouse_NXPE3      PDYNY---------------------------------------HSDTPYYPSG-------         252
Human_NXPE3      AAESA-FFQSGVNIKMPVNSSGPDWVTVIP--RRIKETNSLELSQGSGTFPSGYYYKDQW         331
Human_NXPE4      KQEKSLFERSNVGVEIMEKFNTISVSKCNKETVAMKEKCK---FGMTSTIPSGHVWRNTW         325
Mouse_NXPE4      KQEKSLFERSNVGVEIMEKFNTISVSKCNKETVAMKEKCK---FGMTSTIPSGHVWRNTW         325
Mouse_NXPE2      LEEKLLFRRFNIGVEVVKNLSIVVSLCNNKT-NKKKKKCQ---IGMETPSPGGYTLKGRW         335
Human_NXPE1      DKENSLFHRSKVGVEMMKDRKHIDVTNCNKR-EKIEETCQ---VGMKPPVPGGYTLQGKW         325
Human_NXPE2      KEEWRLFHRSNIGVEMMKNFTPIEVIPCNKS-ENIKKNCQ---IGMKTPFPSGYTLKKMW         336
                                                                   *.*

Mouse_NXPE3      ------------------------------------------------------------         252
Human_NXPE3      RPRKFKMRQFNDPDNITECLQRKVVHLFGDSTIRQWFEYLTTFVPDLVEFNLGSPKNVGP         391
Human_NXPE4      NPVSCSLA----TVKMKECLRGKLIYLMGDSTIRQWMEYFKASINTLKSVDLHESGKLQH         381
Mouse_NXPE4      NPVSCSLA----TVKMKECLRGKLIYLMGDSTIRQWMEYFKASINTLKSVDLHESGKLQH         381
Mouse_NXPE2      ITAHCEQNEFRAIKDINNCLTRKLIYLMGDSTLRQWIYYLPKVVKTLKYFDRHGAGPFKT         395
Human_NXPE1      ITTFCNQVQLDT-IKINGCLKGKLIYLLGDSTLRQWIYYFPKVVKTLKFFDLHETGIFKK         384
Human_NXPE2      ITAFCKQIKFNETKNINDCLERKLIYLMGDSTLHQWIYYLQKAVKTLKYFDHHGAGIFKT         396
```

```
Mouse_NXPE3      ------------------------------------------------------------      252
Human_NXPE3      FLAVDQKHNILLKYRCHGPPIRFTTVFS-NELHYVANELNGIVGGKNTVVAIAVWSHFST      450
Human_NXPE4      QLAVDLDRNINIQWQKYCYPLIGSMTYSVKEMEYLTRAIDRTGGEKNTVIVISLGQHFRP      441
Mouse_NXPE4      QLAVDLDRNINIQWQKYCYPLIGSMTYSVKEMEYLTRAIDRTGGEKNTVIVISLGQHFRP      441
Mouse_NXPE2      HILLDTERHIFVQWKKHSHPFVTNKLFSMKDDNYIPREIDQVAGDSGTAIVISFGQHFRP      455
Human_NXPE1      HLLLDAERHTQIQWKKHSYPFVTFQLYSLIDHDYIPREIDRLSGDKNTAIVITFGQHFRP      444
Human_NXPE2      HVLLDVERHILIQWKKHGHPFVTKKLFSVKDENYIPREIDQVAGDKNTAIVITLGQHFRP      456

Mouse_NXPE3      ------------------------------------------------------------      252
Human_NXPE3      FPLEVYIRRLRNIRRAVVRLLDRSPKTVVVIRTANAQELGPEVSLFNSDWYNFQLDTILR      510
Human_NXPE4      FPIDVFIRRALNVHKAIQHLLLRSPDTMVIIKTENIREMYNDAERF-SDFHGYIQYLIIK      500
Mouse_NXPE4      FPIDVFIRRALNVHKAIQHLLLRSPDTMVIIKTENIREMYNDAERF-SDFHGYIQYLIIK      500
Mouse_NXPE2      FPINVFIRRAINVKNAIERLFLRSPETKVIIKTENIREINEHVEIF-SDFHGSIQNLIIR      514
Human_NXPE1      FPIDIFIRRAIGVQKAIERLFLRSPATKVIIKTENIREMHIETERF-GDFHGYIHYLIMK      503
Human_NXPE2      FPINIFIRRAINIQKAIERLFLRSPETKVILKTENTREIEQNAEMF-SDFHGYIQNLIIR      515

Mouse_NXPE3      ------------------------------------------------- 252
Human_NXPE3      RMFSGVGVYLVDAWEMTLAHYLPHKLHPDEVIVKNQLDMFLSFVCPLET 559
Human_NXPE4      DIFQDLSVSIIDAWDITIA-YGTNNVHPPQHVVGNQINILLNYIC---- 544
Mouse_NXPE4      DIFQDLSVSIIDAWDITIA-YGTNNVHPPQHVVGNQINILLNYIC---- 544
Mouse_NXPE2      DIFRDLNVGIIDAWDMTVA-YRSEDVHPPESVIESQIGMFLNYIC---- 558
Human_NXPE1      DIFKDLNVGIIDAWDMTIA-YGTDTIHPPDHVIGNQINMFLNYIC---- 547
Human_NXPE2      DIFVDLNVGIIDAWDMTIA-YCTNNAHPPDYVIQNQIGMFLNYIC---- 559
```

**Supplementary figure 4. Comparison of NXPE1 with homologue proteins**

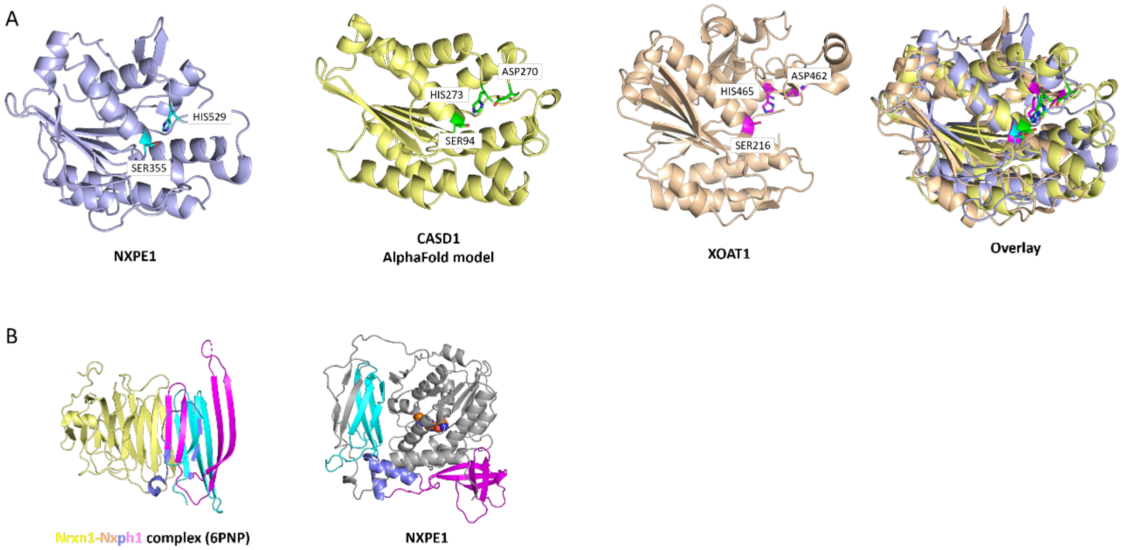


**A**. Comparison and superposition of the esterase domains from NXEP1 (Cryo-EM, light blue), CASD1 (AlphaFold model) [53], and XOAT1 (PDB: 6CCI)[33]. **B**. Comparison of the neurexophilin domain from neurexophilin-1 (PDB 6PNQ)[38] and NXPE1(Cryo-EM). Different elements of neurexcophilin domains are labeled in cyan, blue and magenta in both structures.

**Supplementary figure 5. UPCL chromatographs of wild type and Nxpe2/Nxpe4 dKO Sialic acid glycan species profile**

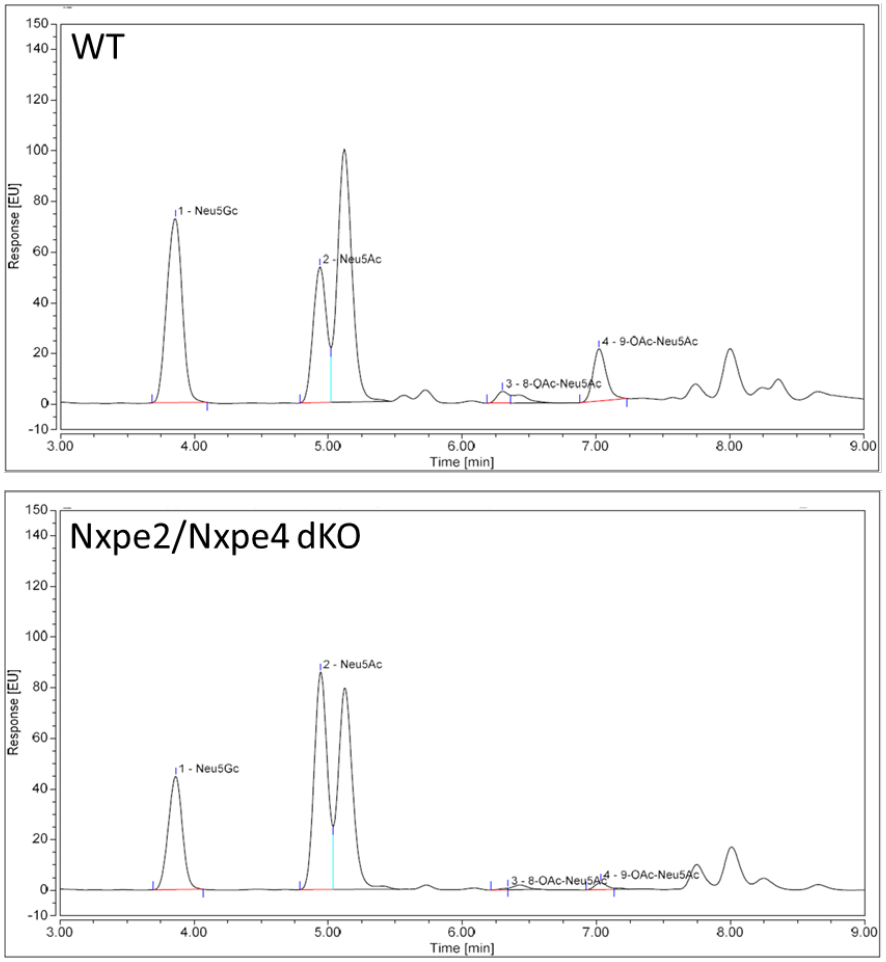
